# A Novel ‘Three-in-one’ Mucosal Vaccine Elicits Broad Protection against Three Distinct Clusters of ACE2-using Sarbecoviruseses

**DOI:** 10.1101/2025.06.26.661659

**Authors:** Lin Liu, Haofeng Lin, Min Li, Xian Li, Shasha Wang, Mengxin Xu, Shuning Liu, Yunqi Hu, Mei-Qin Liu, Zhixiang Huang, Zhen Zhang, Ke Lan, Yu Chen, Huimin Yan, Li Zhou, Jingyi Yang, Yao-Qing Chen, Qingguo Wu

## Abstract

With the persistent risk of coronavirus epidemics, the development of universal vaccines is urgently needed and challenging to achieve. In this study, we aimed to design a multivalent immunogen to provide broad protection against diverse ACE2-using sarbecoviruses, which pose a significant threat to global health. Our analysis revealed that the receptor-binding domains (RBDs) of ACE2-using sarbecoviruses segregate into three antigenically distinct clusters, including one cluster that encompasses viruses related to SARS-CoV-2. Based on these findings, we engineered a ‘three-in-one’ immunogen, designated as 3Rs-NC, which incorporates representative RBDs from all three clusters. 3Rs-NC preserved the natural epitope configuration of each monomeric RBD component, and efficiently elicited high levels of neutralizing antibodies against representative viruses and closely related sarbecoviruses. When administered intranasally with flagellin-based mucosal adjuvant KFD, 3Rs-NC induced robust and durable RBD-specific serum IgG and mucosal IgA responses in mice. Furthermore, KFD-adjuvanted 3Rs-NC conferred sustained protection in both the upper and lower respiratory tracts against SARS-CoV-2 Omicron BA.1 and SARS-like coronavirus WIV1. Additionally, 3Rs-NC immunization protected mice from lethal challenge of SARS-like coronavirus rRsSHC014S, with more efficient protection observed in female mice than male mice. This needle-free ‘three-in-one’ vaccine represents a promising candidate for a universal mucosal vaccine against ACE2-using sarbecoviruses and could serve as a foundational component of ‘three-in-one’ vaccine series to form a comprehensive coronavirus vaccine package.

**Highlight:**

1. novel ‘three-in-one’ immunogen 3Rs-NC induced a high level of neutralizing antibodies against three distinct ACE2-using Sabecovirus clusters.
2. administration of 3Rs-NC plus KFD adjuvant (*3Rs-NC+KFDi.n)* generated potent and long-lasting RBD-specific serum IgG and mucosal IgA.
3. *3Rs-NC+KFDi.n* provided long-term protection in both upper- and lower-respiratory tracts in mice.
4. *3Rs-NC+KFDi.n* could serve as a foundational component for next-generation coronavirus vaccine strategies.

## INTRODUCTION

The COVID-19 pandemic has caused overwhelming hospitalizations, millions of deaths, and trillions of dollars in global economic loss. Importantly, many sarbecoviruses closely related to SARS-CoV-1 and SARS-CoV-2 exist in bats and other mammalian species, serving as potential reservoirs or bridging hosts [1]. However, neutralizing antibody responses elicited by prior SARS-CoV-2 infection or vaccination exhibit limited efficacy against novel sarbecoviruses [2], while the increasing frequency of pathogen emergence from animal reservoirs heightens the risk of future coronavirus pandemics[3]. Consequently, the World Health Organization (WHO) has identified coronavirus as one of the high-priority pathogens for a potential future pandemic.

To combat this ongoing threat, developing next-generation coronavirus vaccines remains a public health priority [4]. The SARS-CoV-2 pandemic revealed three critical requirements for ideal vaccines against coronaviruses. First, effective vaccines should induce robust mucosal immunity in the upper respiratory tract where initial infection predominantly occurs [5], in addition to systemic immunity, to prevent the establishment of infection rather than merely reducing disease symptoms [18]. Second, long-lasting protective immunity, especially neutralizing antibodies, is required to overcome short incubation periods of coronaviruses, which limit memory recall efficacy [4]. Third, the achievement of high-level cross-protection against diverse coronavirus variants is imperative[6], although developing pan-genus or pan-family vaccines remains challenging due to the difficulty in generating broad-spectrum neutralizing antibodies [7]. Given these challenges, we propose an alternative strategy that is to develop a modular package of vaccines covering major coronaviruses. Each mucosal vaccine in this package could induce high-level and long-lasting neutralizing antibodies, with efficient manufacturing. In the event of a coronavirus epidemic, a suitable vaccine candidate could be immediately selected from this vaccine package, validated by rapid neutralization assays of pre-stored immune sera.

In previous studies, we found that intranasal (i.n) inoculation of recombinant triple receptor binding domain (RBD) proteins with a flagellin-based mucosal adjuvant could induce long-lasting neutralizing antibodies in serum and mucosal sites, offering durable protection in both the upper- and lower-respiratory tracts [8, 9]. Notably, this approach also elicited neutralizing antibody responses and mucosal IgA responses in human volunteers with minimal adverse effects [9]. We hypothesized that a similar strategy could be used to develop mucosal vaccines against targeted coronaviruses. Prioritizing high-threat coronaviruses, we focused on generating a mucosal vaccine against human angiotensin-converting enzyme 2 (ACE2)-using sarbecoviruses. Notably, sarbecoviruses exhibit substantial genetic diversity in their RBDs that determine host range, and vary in affinity for the human ACE2 receptor [10, 11]. Human ACE2-using sarbecoviruses pose a significant threat due to their capacity for human ACE2-dependent entry and replication in human cells [12].

Through RBD sequence analysis, three antigenically distinct clusters of human ACE2-using sarbecoviruses were identified in this study. Among them, SARS-CoV-2 Omicron BA.1, and two SARS-like coronaviruses WIV1 and RsSHC014 isolated from southwest China with a broad species tropism [13], were selected as representative RBD strains of each cluster. Using a ‘three-in-one’ strategy, we designed a recombinant protein 3Rs-NC incorporating the three representative RBDs into a scaffold NC. Compared to a previously reported immunogen 3Ro-NC that only contains RBDs of SARS-CoV-2 [8], 3Rs-NC was superior in terms of eliciting broad-spectrum neutralizing antibodies against various sarbecoviruses. Our results further demonstrate that the novel immunogen 3Rs-NC in combination with a flagellin-based KFD adjuvant induced robust and durable humoral immune responses, providing protection against all three clusters of sarbecoviruses.

## RESULTS

### The three-in-one protein 3Rs-NC preserves native 3D structures of the RBDs

According to the phylogenetic tree of amino acid sequences, the RBDs of sarbecoviruses (**Supplementary Table 1**) could be divided into: clade1a, clade 1b, clade 2, and clade 3 (**Fig.1A**) [14]. Clades 1a and 1b includes ACE2-using sarbecoviruses, containing SARS-CoV-1 and SARS-CoV-2, respectively (**Fig.1A**). Principal coordinate analysis (PCoA) of antigenic distances indicated that the RBDs of clade 1a and 1b could be further divided into three distinct antigenic clusters (**Fig.1B**). The RBDs of clade 1b form a single cluster, while RBDs of clade 1a segregate into two clusters. Notably, the clade 1b virus SARS-CoV-1 clusters with SARS-like virus WIV1, but not with RsSHC014 (**Fig.1B**).

**Fig. 1.**
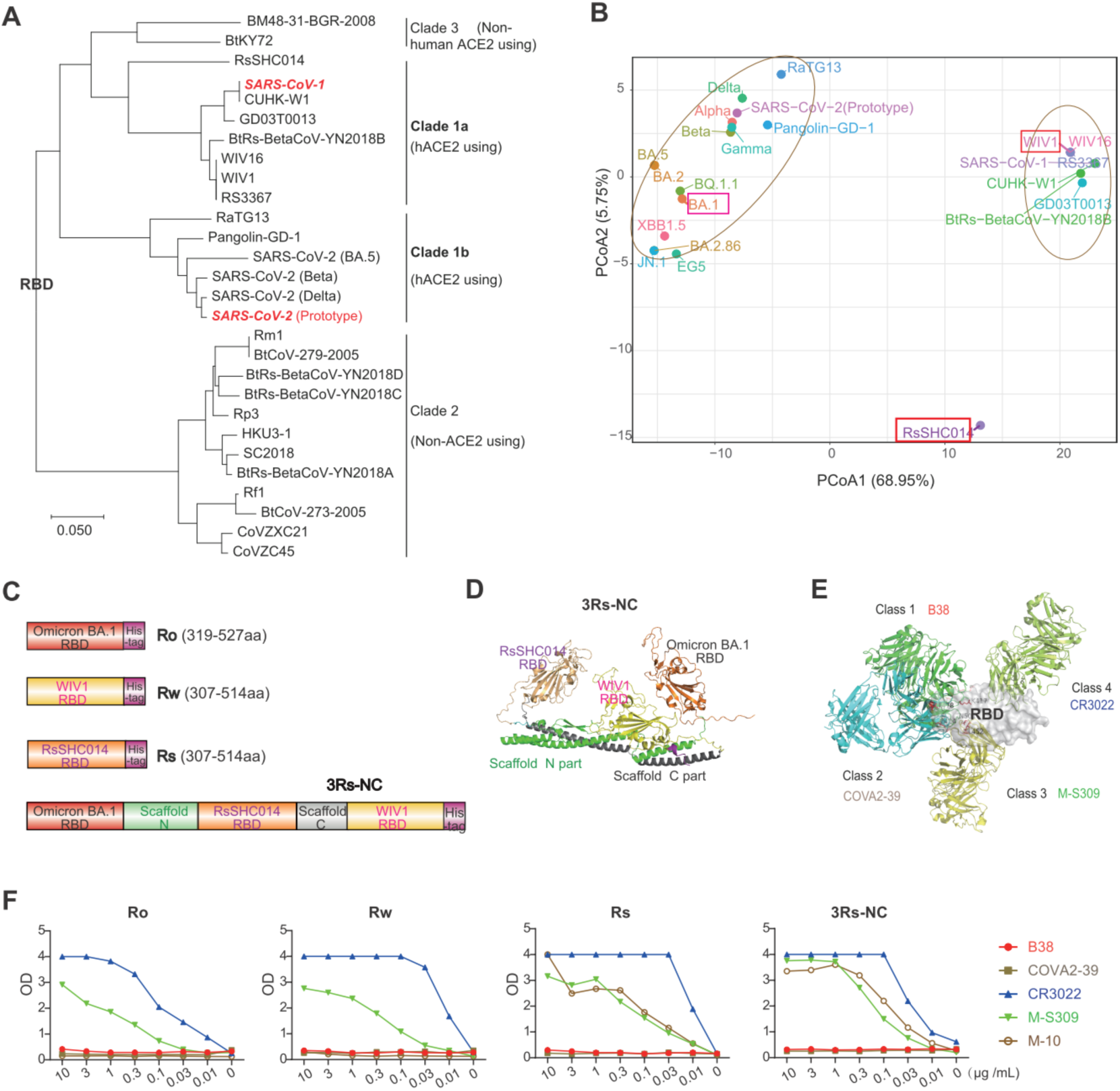
Construction of 3Rs-NC containing RBDs from three distant sarbecovirus. (**A**) Phylogenetic tree of sarbecovirus RBD sequences by maximum likelihood analyses. The scale bar indicates the genetic distance scale of 5%. (**B**) Principal coordinate analysis (PCoA) of antigenic distances among viruses of clade 1a and clade 1b sarbecovirus. The axes represent the two highest variance components. Each dot corresponds to a viral strain as labeled. (**C**) Schematic diagram of the recombinant protein 3Rs-NC containing triple RBDs. RBD genes of SARS-CoV-2 Omicron BA.1 strain, SARS-like virus WIV1, and SARS-like virus RsRSHC014 were connected by the gene of scaffold *NC* to generate *3Rs-NC* gene. (**D**) 3D structure of the protein 3Rs-NC predicted by AlphaFold 3. (**E**) Diagram of the RBD binding with four classes of SARS-CoV-2 RBD-specific neutralizing monoclonal antibodies (mAbs). (**F**) Binding ability of SARS-CoV-2 Omicron BA.1 strain RBD (Ro), WIV1 RBD (Rw), RsRSHC014 RBD (Rs), and 3Rs-NC with the representative neutralizing mAbs analyzed by ELISA. Data shown represent one of two independent experiments each with a duplicate.

To develop a pan-ACE2-using sarbecoviruses vaccine based on the RBD of viruses, we designed a novel immunogen that incorporates RBDs representative of all three antigenic clusters. Given that RBDs of SARS-CoV-2 Omiron BA.1, WIV1, and RsSHC014 are located in these respective clusters, and considering the availability of challenge models for the corresponding viruses [15], these three RBDs were selected as representative immunogens. The three RBDs were then connected by the N-terminal (N) and C-terminal (C) scaffold domains, which have been previously utilized in our multivalent antigen design [8], to generate a ‘Three-in-one’ recombinant construct, named 3Rs-NC (**Fig. 1C** and **D**).

The structure prediction using AlphaFold 3 indicated that the RBD regions of SARS-CoV-2 Omicron subvariant BA.1, WIV1, and RsSHC014 in 3Rs-NC preserved their native 3D structures (**Supplementary** Figure 1). To further test the reactivity and conformation of the RBDs in 3Rs-NC, monoclonal antibodies (mAbs) representing four classes of RBD-specific neutralizing antibodies against SARS-CoV-2 prototype [16] were generated (**Fig. 1E**). Both monomeric RBDs (Omicron variant BA.1 (Ro), WIV1(Rw), RsSHC014 (Rs)) and trivalent 3Rs-NC exhibited vanished binding ability to Class 1 (B38) and Class 2 (COVA2-39) antibodies, which have high neutralizing potency against SARS-CoV-2 prototype (**Fig. 1F**), while maintaining binding affinity to Class 3 (M-S309) and Class 4 (CR3022) antibodies with low neutralizing potency (**Fig. 1F**). Notably, 3Rs-NC showed relatively higher binding affinity than monomeric Rw, Rs, and especially Ro (**Fig. 1F**). 3Rs-NC also showed relatively higher binding affinity than Rs for a RsSHC014 RBD-specific mAb M-10 [17]. These results collectively indicate that all three RBD regions in 3Rs-NC retain their native conformation structures, similar to those in the monomeric RBDs.

### 3Rs-NC immunization elicits potent and broad-spectrum neutralizing antibodies

To assess the immunogenicity of 3Rs-NC, BALB/c mice were intramuscularly immunized three times with a 4 µg/dose of 3Rs-NC or a combination of monomeric RBDs (Ro+Rs+Rw), both formulated with aluminum adjuvant (AL-adjuvant). A previously reported immunogen 3Ro-NC [8], which contains two RBDs of Omicron BA.1 and one RBD of SARS-CoV-2 delta strain within an identical scaffold NC (**Fig. 2A**), was selected for comparison. Saline alone was given as a negative control. (**Fig. 2**B).

**Fig. 2.**
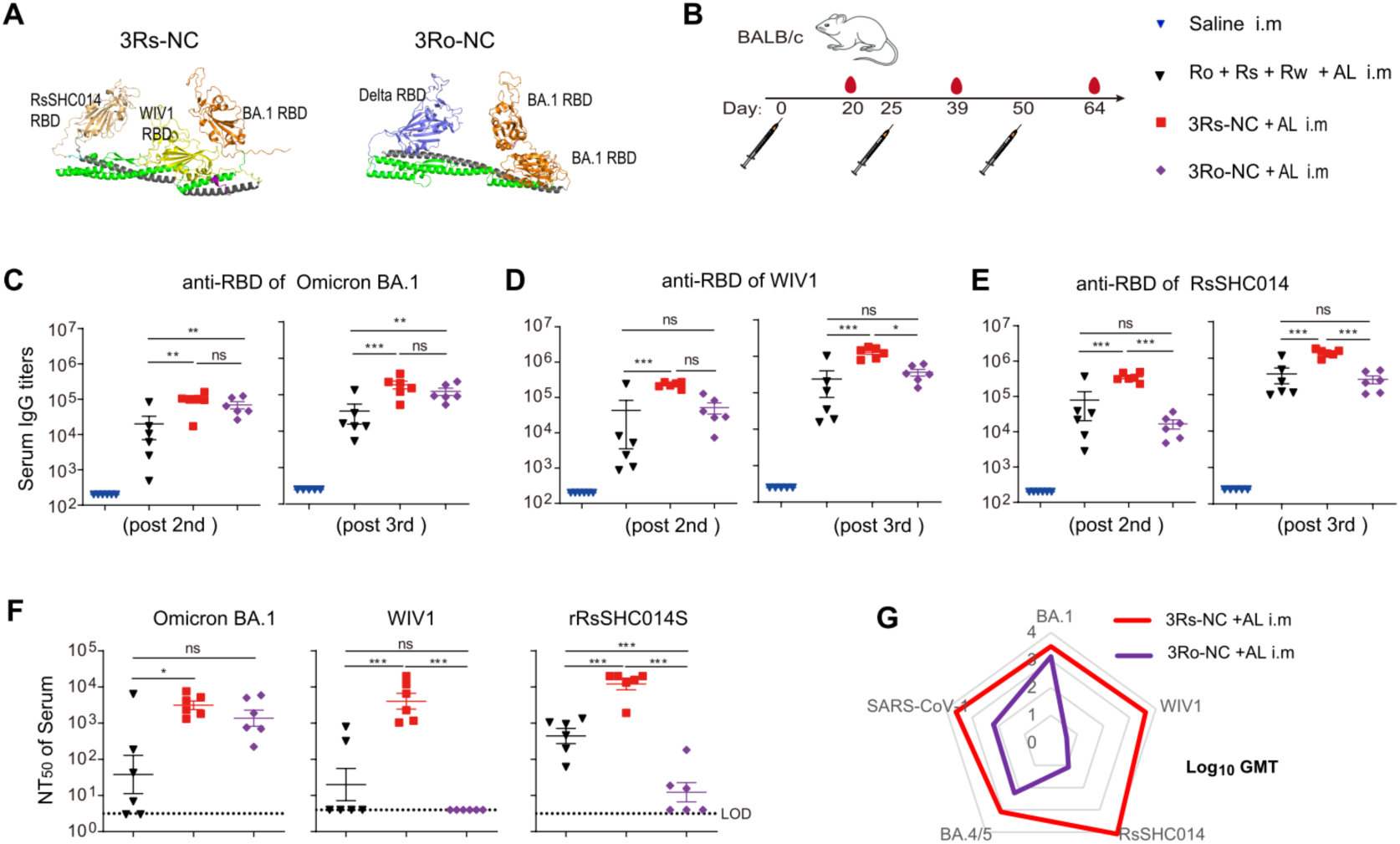
Immunogenicity of 3Rs-NC in the presence of aluminum adjuvant in BALB/c mice. (**A**) The AlphaFold 3 predicted 3D structure of 3Rs-NC, and 3D structure of 3Ro-NC that contains One RBD of SARS-CoV-2 Delta strain and two RBDs of Omicron BA.1 strain in scaffold NC. (**B**) Diagram scheme of immunization and sampling (n=6 female mice per group). (**C-E**) RBD-specific IgG responses in serum after the 2^nd^ and 3^rd^ immunizations. (**F**) Neutralizing antibody responses in serum after the 3^rd^ immunization against virus Omicron BA.1, WIV1, and rRsSHC014S. (**G**) Geometric mean titers (GMT) of neutralizing antibodies against three authentic viruses and two pseudotyped viruses SARS-CoV-1 and SARS-CoV-2 variants BA.4/5. Data are represented as mean ± SEM and are representative of at least two independent experiments. In (C-F), groups were compared using one-way ANOVA. *p < 0.05; **p < 0.01; ***p < 0.001; ns, nonsignificant.

After the 2^nd^ and 3^rd^ immunizations, 3Rs-NC elicited considerably greater RBD-specific IgG responses than Ro+Rs+Rw against all targeted RBDs (**Fig. 2C-E**). Consistently, sera from 3Rs-NC immunized mice exhibited much higher neutralizing ability against authentic viruses SARS-CoV-2 Omicron BA.1, WIV1, and RsSHC014 compared to Ro+Rs+Rw sera (**Fig. 2F**), indicating superior immunogenicity of 3Rs-NC for generating a high level of neutralizing antibodies.

While 3Ro-NC induced SARS-CoV-2 Omicron BA.1 RBD-specific serum IgG and neutralizing antibodies against Omicron BA.1 comparable to those elicited by 3Rs-NC, its neutralizing antibodies against WIV1 and rRsSHC014S were nearly undetectable, with titers more than 200-fold lower than those in 3Rs-NC (**Fig. 2C-F**). Compared to Ro+Rs+Rw, 3Ro-NC generated comparable RsSHC014 RBD-specific binding antibodies after the 3^rd^ immunization but more than 20-fold lower neutralizing antibodies against rRsSHC014S (**Fig. 2E** and **F**). These suggest that while cross-reactive binding antibodies to RBDs are readily generated, neutralizing antibodies are poorly elicited by distantly related RBDs. Both 3Rs-NC and Ro+Rs+Rw induced stronger antibody responses against WIV1 and RsSHC014 than against SARS-CoV-2 Omicron BA.1, in both binding and neutralizing potency (**Fig. 2C-F**), indicating lower immunogenicity of Omicron BA.1 RBD than that of WIV1 and RsSHC014 RBDs.

Pseudovirus neutralization assay further revealed that 3Rs-NC induced higher levels of neutralizing antibody responses against SARS-CoV-2 Omiron BA.4/5 and SARS-CoV-1 than 3Ro-NC (**Supplementary Figure 2**). The geometric mean titer (GMT) of neutralizing antibodies against the sarbecoviruses indicated that 3Rs-NC generated a broader spectrum of neutralizing antibodies than 3Ro-NC (**Fig. 2G**). These results suggest that the combination of RBDs from antigenically distant sarbecoviruses with the NC scaffold represents a promising strategy for eliciting potent and broad-spectrum neutralizing antibody responses.

### Intranasal immunization with adjuvanted 3Rs-NC elicits coordinated systemic and mucosal immunity

Developing effective subunit mucosal vaccines requires potent mucosal adjuvants. In this study, we selected CpG and the flagellin-derived recombinant protein KFD [18–21] as mucosal adjuvants. Briefly, BALB/c mice were intranasally immunized three times with either 4 µg of 3Rs-NC alone (*3Rs-NCi.n*), 4 µg of 3Rs-NC plus 1 µg of KFD adjuvant (*3Rs-NC+KFDi.n*), or 4 µg of 3Rs-NC plus 1 µg of CpG ODN 2395 adjuvant (*3Rs-NC+CpGi.n*). Control groups received either saline alone or intramuscular immunization with 4 µg of 3Rs-NC plus 200 µg of AL-adjuvant (*3Rs-NC+ALi.m*) (**Fig. 3A**). Mouse serum and mucosal samples were collected at 14 days after the third immunization to assess the humoral responses.

**Fig. 3.**
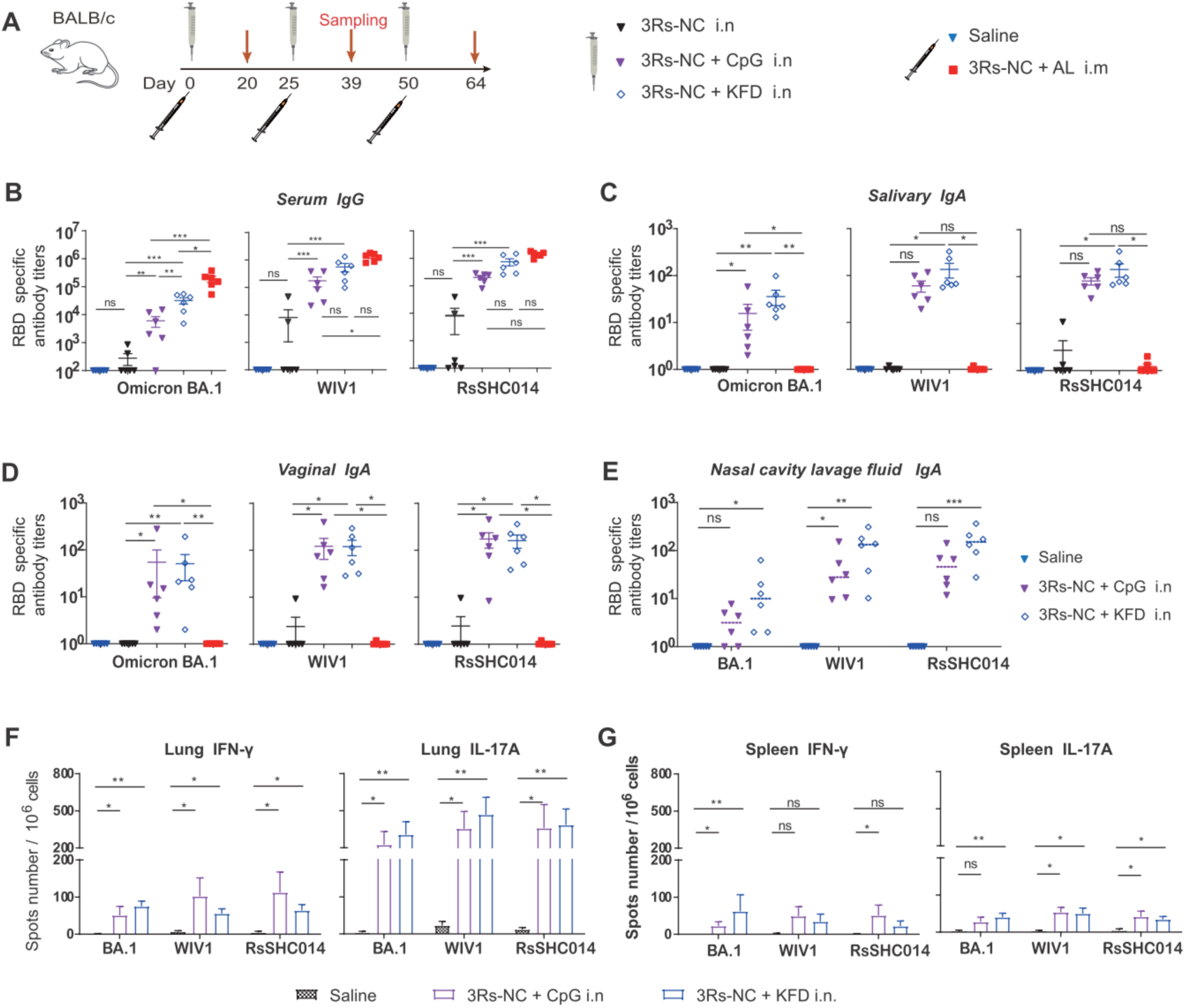
Antibody and T cell responses induced by intranasal immunization of 3Rs-NC adjuvanted with KFD or CpG in BALB/c mice. (**A**) Diagram scheme of immunization and sampling (n=6 female mice per group). (**B**) RBD-specific IgG responses in serum after the 3^rd^ immunization. (**C-E**) RBD-specific mucosal IgA responses after the 3^rd^ immunization in saliva (**C**), vaginal lavage fluid (**D**), and nasal turbinate lavage fluid (**E**). (**F** and **G**) RBD-specific T cell responses after the 3^rd^ immunization in lung (**F**) and spleen (**G**). Data are represented as mean ± SEM and are representative of at least two independent experiments. Groups were compared using one-way ANOVA. *p < 0.05; **p < 0.01; ***p < 0.001; ns, nonsignificant.

The results showed that intranasal immunization with 3Rs-NC elicited RBD-specific serum IgG, salivary IgA, and vaginal IgA in mice, only in the presence of mucosal adjuvants (**Fig. 3B-D**). As expected, antigen-specific mucosal IgA antibodies could be induced by the intranasal immunization route, but not by intramuscular immunization. (**Fig. 3C-E**). *3Rs-NC+KFDi.n* induced slightly lower serum IgG than *3Rs-NC+ALi.m*, but the difference was non-significant except for the SARS-CoV-2 Omicron BA.1 RBD-specific response (**Fig. 3B**). Notably, compared to CpG-adjuvanted immunogen, KFD-adjuvanted 3Rs-NC induced a higher level of RBD-specific serum IgG (**Fig. 3B**), salivary IgA (**Fig. 3C**), and nasal IgA responses (**Fig. 3E**), particularly in eliciting Omicron BA.1 RBD-specific antibody responses (**Fig. 3B-D**). These results indicated that KFD exhibited greater adjuvant activity than CpG for intranasally administered 3Rs-NC.

We further evaluated the immunogenicity of varying doses of immunogen when combined with 1 µg of KFD as a mucosal adjuvant. Dose-response analysis revealed that intranasal inoculation with 4 µg of 3Rs-NC induced significantly higher levels of RBD-specific antibody responses than inoculation with 1.3 µg of 3Rs-NC, but comparable levels to inoculation with 12 µg of 3Rs-NC (**Supplementary** Figure 3). Collectively, 3Rs-NC is highly immunogenic when adjuvanted with KFD through intranasal immunization, eliciting coordinated systemic and mucosal immunity against Omicron BA.1, WIV1, and RsSHC014.

Analysis of T cell responses revealed that both *3Rs-NC+KFDi.n* and *3Rs-NC+CpGi.n* induced robust RBD-specific IFN-γ (Th1) and IL-17A (Th17) secreting T cell responses, both in the lung and in the spleen (**Fig. 3F** and **G**). Notably, in the lung tissue, the RBD-specific Th17 responses were much higher than RBD-specific Th1 responses, suggesting a significant bias towards Th17 responses in the lung for both *3Rs-NC+KFDi.n* and *3Rs-NC+CpGi.n*.

### *3Rs-NC+KFDi.n* protects the upper and lower respiratory tracts against infection

To further investigate the protective efficacy of the 3Rs-NC vaccine, HFH4-human ACE2 transgenic mice (hACE2) were utilized to evaluate the immune response and protection against the infection [22]. In brief, mice were immunized with 4 µg of 3Rs-NC plus 1 µg of KFD via the intranasal route (*3Rs-NC+KFDi.n*), or with 3Rs-NC plus AL-adjuvant via the intramuscular route (*3Rs-NC+ALi.m*), using saline inoculation as the control group. After three doses of immunization, each group of mice was divided into the SARS-CoV-2 Omicron BA.1 subgroup and the WIV1 subgroup, which were challenged with SARS-CoV-2 Omicron variant BA.1 and WIV1, respectively (**Fig. 4A**).

**Fig. 4.**
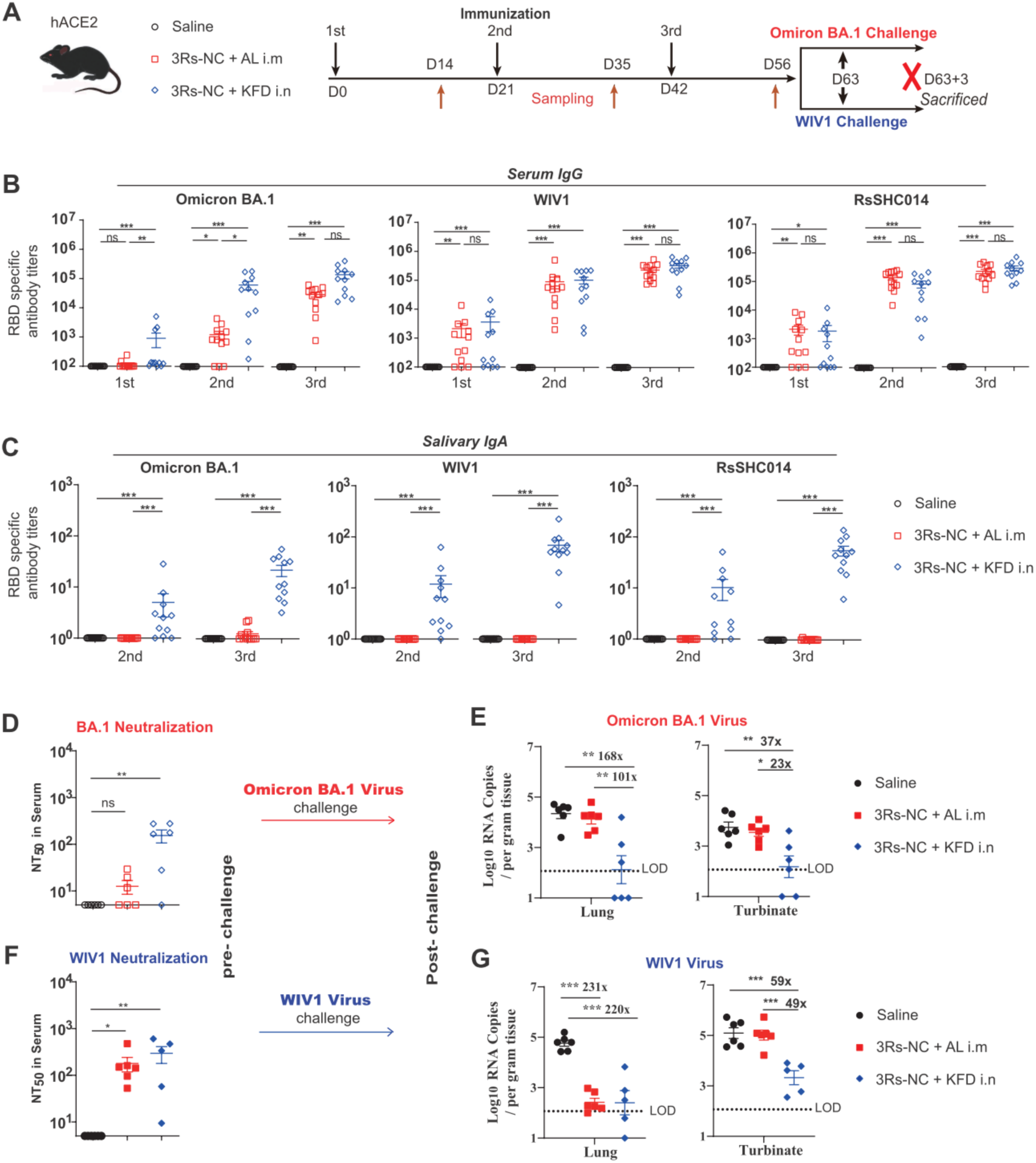
Protection of 3Rs-NC immunization against SARS-CoV-2 Omicron BA.1 and WIV1 infection in female HFH4-hACE2 mice. (**A**) Diagram scheme of immunization and virus challenge. Each group (n = 12) of mice was divided into two subgroups after immunization and challenged by Omicron BA.1 or WIV1, respectively. (**B**) RBD-specific serum IgG after the 1^st^, 2^nd^ and 3^rd^ immunizations. (**C**) RBD-specific salivary IgA after the 2^nd^ and 3^rd^ immunizations. (**D** and **F**) Neutralization antibody responses against authentic SARS-CoV-2 Omicron BA.1 in serum after the 3^rd^ immunization and ahead of the challenge. (**E** and **G**) qPCR tested RNA copies of SARS-CoV-2 RBD in lung and turbinate at 3 days post-infection. Groups were compared using one-way ANOVA. *p < 0.05; **p < 0.01; ***p < 0.001; ns, nonsignificant. LOD, limit of detection.

Consistent with the immune responses observed in BALB/c mice, both *3Rs-NC+KFDi.n* and *3Rs-NC+ALi.m* induced robust serum IgG antibody responses against the RBDs of Omicron BA.1, WIV1, and RsSHC014 in hACE2 mice (**Fig. 4B**). Notably, only intranasal immunization (*3Rs-NC+KFDi.n*) generated RBD-specific IgA in saliva (**Fig. 4C**) and vaginal lavage fluid (Supplementary Figure 4).

Different from the findings in BALB/c mice, *3Rs-NC+KFDi.n* showed comparable efficiency to *3Rs-NC+ALi.m* in inducing WIV1 and RsSHC014 RBD-specific serum IgG responses in hACE2 mice (**Fig. 4B**). Moreover, for the induction of SARS-CoV-2 Omicron BA.1 RBD-specific serum IgG responses, *3Rs-NC+KFDi.n* exhibited significantly higher efficiency than *3Rs-NC+ALi.m* in hACE2 mice, particularly after the 1^st^ and 2^nd^ immunizations (**Fig. 4B**). The results of neutralization assay further revealed that *3Rs-NC+KFDi.n* immunization elicited higher levels of neutralizing antibodies against authentic SARS-CoV-2 Omicron BA.1, and comparable neutralizing antibody titers against authentic WIV1, compared to the *3Rs-NC+ALi.m* group (**Fig. 4D, E**).

At 21 days after the third immunization, mice were challenged with either SARS-CoV-2 Omicron BA.1 or WIV. At 3 days post-infection (dpi), viral loads in lungs and nasal turbinate tissues were determined using RT-qPCR. In Omicron BA.1-challenged mice, viral genome copies in the lungs of *3Rs-NC+KFDi.n* immunized group were significantly reduced by 168-fold compared to the saline control group and by 101-fold compared to the *3Rs-NC+ALi.m* group (**Fig. 4E**, left panel). In the *3Rs-NC+ALi.m* group, the reduction in Omicron viral load in lung tissue was not significant (**Fig. 4E**, left panel), consistent with the low levels of Omicron BA.1-specific antibody response in this group (**Fig. 4B** and **D**). In nasal turbinate tissues, only the *3Rs-NC+KFDi.n* group conferred protection with 37-fold and 23-fold viral RNA reductions, compared to the saline group and the *3Rs-NC+ALi.m* group, respectively (**Fig. 4E**, right panel).

In WIV1-challenged mice, immunization with both *3Rs-NC+KFDi.n* and *3Rs-NC+ALi.m* significantly reduced viral genome copies in lung tissue compared to the saline control group (**Fig. 4G**, left panel, 220-fold and 231-fold respectively). In the nasal turbinate, only the *3Rs-NC+KFDi.n* group showed a substantial decrease in viral load, with reductions of 59-fold compared to the saline group and 49-fold compared to the *3Rs-NC+ALi.m* group (**Fig. 4G**, right panel).

These results indicate that intranasal immunization with 3Rs-NC plus KFD adjuvant can provide protection against infection in both upper- and lower-respiratory tracts as a prophylactic mucosal vaccine against SARS-CoV-2 omicron and WIV1.

### *3Rs-NC+KFDi.n* elicits enhanced protection against rRsSHC014S in female mice

Male HFH4-hACE2 mice have been reported to be more susceptible to lethal outcomes following sarbecoviruses challenge [22]. To evaluate whether *3Rs-NC+KFDi.n* could induce robust RBD-specific antibody responses and offers protection against lethal sarbecoviruses challenge in both female and male mice, HFH4-hACE2 mice were immunized with *3Rs-NC+KFDi.n* or *3Rs-NC+ALi.m*, using saline inoculation as control. After three doses of immunization, mice were challenged with a lethal dose of rRsSHC014S (**Fig. 5A**).

**Fig. 5.**
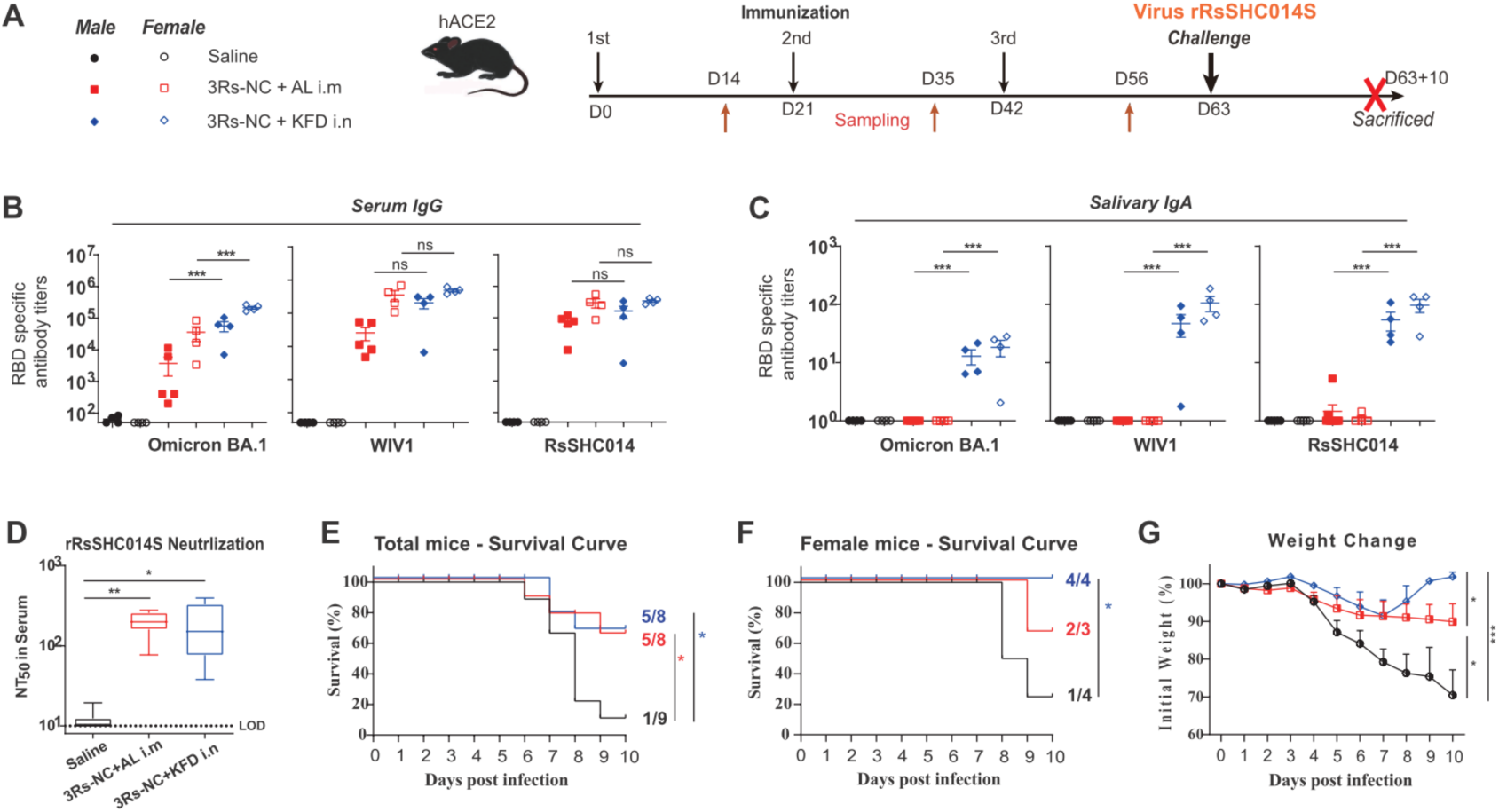
Protection of 3Rs-NC immunization against virus rRsSHC014S in male and female HFH4-hACE2 mice. (**A**) Diagram scheme of immunization and virus challenge (n = 8-9, including 4 female mice). (**B** and **C**) RBD-specific serum IgG (**B**) and salivary IgA (**C**) after the 3^rd^ immunization. (**D**) Neutralization antibody responses against rRsSHC014S in serum after the 3^rd^ immunization and ahead of the challenge. (**E** and **F**) Survival curves of total mice (**E**) and female mice (**F**) within 10 days post-infection. (**G**) Weight changes of total mice within 10 days post-infection. Groups were compared using one-way ANOVA. *p < 0.05; **p < 0.01; ***p < 0.001; ns, nonsignificant.

In both female and male hACE2 mice, 3Rs-NC immunization induced strong antigen-specific serum IgG antibodies, although IgG responses in male mice were slightly weaker than those observed in female mice (**Fig. 5B** and **supplementary figure 5**). RBD-specific IgA in saliva was also induced in mice immunized with *3Rs-NC+KFDi.n*, regardless of sex (**Fig. 5C**). In line with the RBD-specific serum IgG response, sera from *3Rs-NC+KFDi.n* immunized mice exhibited neutralizing potential against rRsSHC014S virus, similar to that of *3Rs-NC+ALi.m* (**Fig. 5D**).

After rRsSHC014S virus challenge, 8 of 9 mice in the saline control group succumbed (or were sacrificed after losing above 30% of body weight) within 10 days post-infection (**Fig. 5E**). In contrast, only 3 of 8 mice died post-challenge in both *3Rs-NC+KFDi.n* group and *3Rs-NC+ALi.m* group (**Fig. 5E**). These findings indicate that both *3Rs-NC+KFDi.n* and *3Rs-NC+Ali.m* immunization could offer protection against the lethal challenge of rRsSHC014S.

Further analysis of sex-specific protection revealed distinct levels of protection efficacy in female and male mice. Among female mice, all survived in the *3Rs-NC+KFDi.n* group, whereas 3 of 4 female mice died in the saline control group (**Fig. 5F**). Among male mice, 3 deaths occurred in 4 mice of the *3Rs-NC+KFDi.n* group, while all 5 mice died in the saline control group (**Supplementary Figure 6A**). Weight change curves showed that body weight reduction in the *3Rs-NC+KFDi.n* group was less than that in both the saline group and the *3Rs-NC+ALi.m* group (**Fig. 5G**). In female mice of the *3Rs-NC+KFDi.n* group, weight loss was nearly undetectable after rRsSHC014S challenge (**Fig. Supplementary Figure 6B**). These results suggest that protection conferred by *3Rs-NC+KFDi.n* was more prominent in female mice than male mice, indicating superior efficacy in females.

### *3Rs-NC+KFDi.n* induces long-lasting RBD-specific antibodies and confers long-term protection

To determine whether the protective antibody responses could be long-lasting, we monitored immune responses for up to 12 months following the 3^rd^ immunization, and challenged mice with the BALB/c mice-sensitive Omicron BA.1 strain at 12.5 months after the 3^rd^ immunization (**Fig. 6A**).

**Fig. 6.**
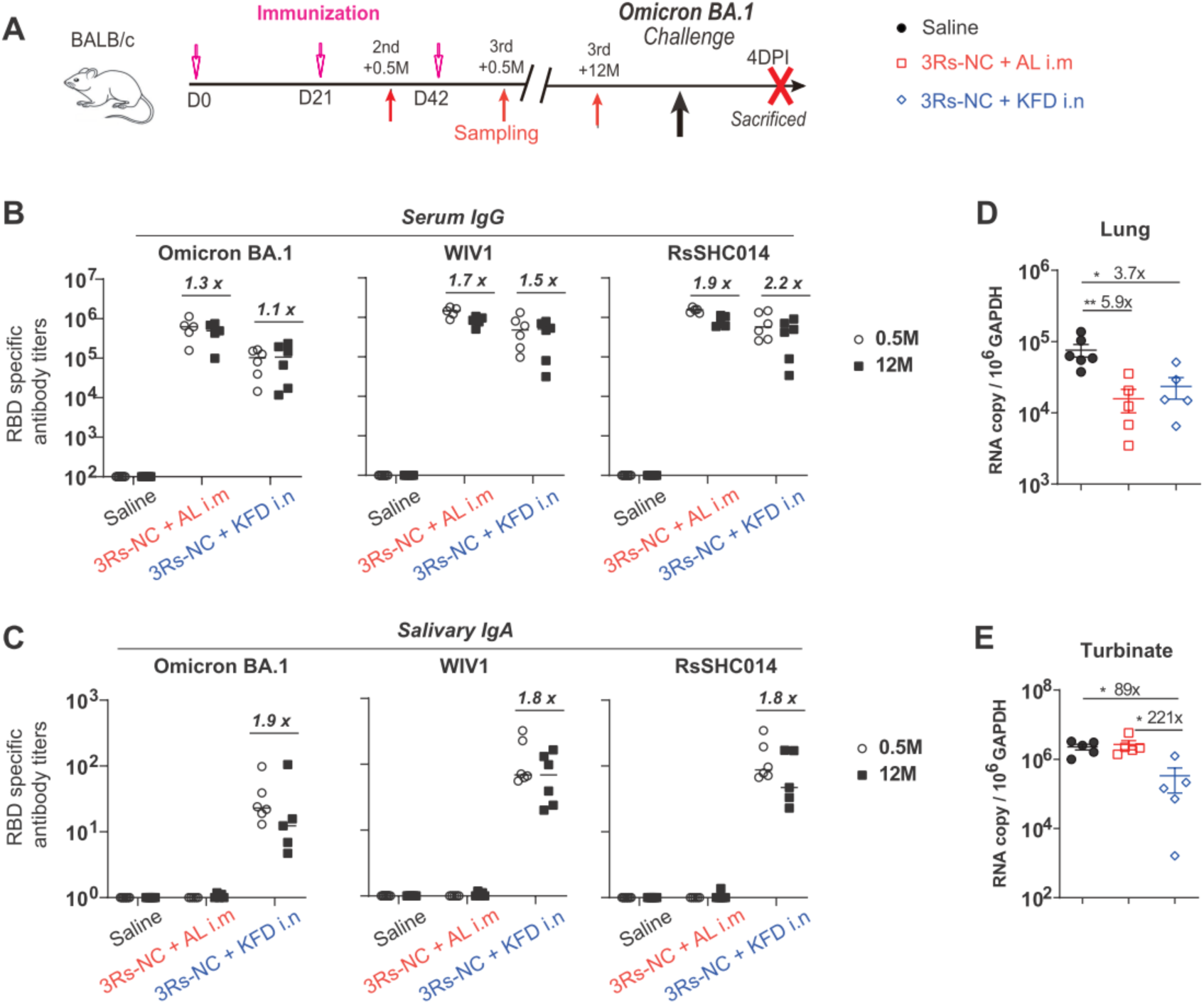
Long-lasting antibody responses and protection of 3Rs-NC immunization against SARS-CoV-2 Omicron BA.1. (**A**) Diagram scheme of immunization and virus challenge (n = 5-6 female BALB/c mice per group). (**B** and **C**) RBD-specific serum IgG and salivary IgA at 0.5 months (0.5M) and 12 months (12M) after the 3^rd^ immunization. (**D** and **E**) qPCR tested RNA copies of SARS-CoV-2 E gene in lung and turbinate at 4 days post-infection. Groups were compared using one-way ANOVA. *p < 0.05; **p < 0.01.

RBD-specific IgG against Omicron BA.1, WIV1, and RsSHC014 showed no significant reduction at 12 months after the 3^rd^ immunization compared to levels at 0.5 months after the 3^rd^ immunization (**Fig. 6B**). Furthermore, RBD-specific IgA responses detected in *3R-NC+KFDi.n* group in saliva (**Fig. 6C**) and in virginal lavage fluid (**Supplementary** Figure 7) persisted long-term, with less than 2-fold reduction within 11.5 months. These results demonstrate that both *3Rs-NC+KFDi.n* and *3Rs-NC+ALi.m* induce durable RBD-specific antibody responses.

Following challenge with the Omicron BA.1 strain, copy numbers of viral *E* gene and internal reference *GAPDH* gene in lungs and in turbinate tissues were evaluated at 4dpi. In the lungs, compared to the saline immunized group, both the *3Rs-NC+KFDi.n* group and the *3Rs-NC+ALi.m* group of mice showed significant reductions in virus infection (**Fig. 6D**, 3.7-fold and 5.9-fold respectively). In turbinate tissues, compared to the saline immunized group, only the *3Rs-NC+KFDi.n* group of mice showed a significant reduction in virus infection (**Fig. 6E**, 89-fold). Furthermore, compared to the *3Rs-NC+ALi.m* immunized mice, the *3Rs-NC+KFDi.n* group of mice also exhibited a significant reduction in virus infection in turbinate tissues (**Fig. 6E**, 221-fold). These results demonstrate that *3Rs-NC+KFDi.n* confer long-term protection against SARS-CoV-2 variant infection in both upper- and lower-respiratory tracts.

## DISCUSSION

In the development of pan-coronavirus vaccines, spike protein-derived antigens, particularly the receptor binding domain (RBD), are widely selected as immunogenic components due to the high efficacy in inducing neutralizing antibodies [23, 24] and the ease of manufacturing [25]. Several approaches have been attempted to generate cross-neutralizing coronavirus antibodies, with one effective method involving the covalent attachment of up to eight different coronavirus RBDs to nanoparticles to create mosaic RBD-NPs [26, 27]. However, achieving broad-spectrum protection against emerging coronavirus variants remains a significant challenge in vaccinology [7].

To address this limitation, we propose an alternative strategy: developing a modular vaccine package where each component is designed to induce high levels of neutralizing antibodies against several specific coronavirus subgroups. As a priority, we focused on generating a universal mucosal vaccine against all clusters of human ACE2-using sarbecoviruses, which present a high threat potential [1].

Analysis of RBD sequences revealed three distinct clusters within human ACE2-using sarbecoviruses (Fig. 1). From each cluster, we selected one representative strain, SARS-CoV-2 Omiron BA.1, WIV1, and RsSHC014 to engineer a ‘three-in-one’ trivalent immunogen 3Rs-NC. This novel immunogen exhibited significantly enhanced immunogenicity compared to a mixture of monomeric RBDs (Fig. 2). In addition, unlike 3Ro-NC, which contains two RBDs of SARS-CoV-2 Omicron BA.1strain and one RBD of Delta strain, 3Rs-NC induced robust neutralizing antibody responses against SARS-like coronavirus WIV1 and rRsSHC014S (Fig. 2). This aligns with previous reports indicating that RBDs derived from SARS-CoV-2 strains are less effective in inducing neutralizing antibodies against WIV1 and rRsSHC014S [26, 28].

Notably, in terms of inducing RBD-specific serum IgG responses, although *3Rs-NC+KFDi.n* induced moderately lower serum IgG than *3Rs-NC+ALi.m* in BALB/c mice (**Fig. 3B**), it generated comparable responses against WIV1 and RsSHC014 (**Fig. 4B**) and superior responses against the SARS-CoV-2 Omicron strain in hACE2 mice, particularly after the 1^st^ and 2^nd^ immunization (**Fig. 4B**). Whether these differences in humoral immune responses are related to hACE2 expression warrants further investigation.

As a prophylactic mucosal vaccine*, 3Rs-NC+KFDi.n* provided protection against SARS-CoV-2 omicron and SARS-like WIV1 infections in both upper- and lower-respiratory tracts (**Fig. 4**). Consistent with previous studies [8, 29], the reduction of viral load in lung and turbinate tissues correlated with neutralizing antibody levels in serum and RBD-specific IgA antibodies in the upper respiratory tract, respectively. In addition, RBD-specific IgG and mucosal IgA generated by *3Rs-NC+KFDi.n* are long-lasting, offering sustained protection in both upper- and lower-respiratory tracts for at least 12 months (**Fig. 6**).

T-cell responses, especially tissue-resident memory T cells in the respiratory tract, play a crucial role in pan-sarbecovirus protective immunity [1, 30]. Research on adenoviral-vectored vaccines has shown that RBD-specific CD4^+^ and CD8^+^ T-cell-mediated anamnestic responses could confer protection across sarbecovirus clades [30]. Additionally, previous studies on COVID-19 patients also highlight that the presence and magnitude of T-cell responses were both positively correlated with protection against SARS-CoV-2 infection [31, 32]. In this study, we observed that *3Rs-NC+KFDi.n* induced RBD-specific Th1 and Th17 responses in spleen and lung tissues (**Fig. 4**). In addition to IFN-γ-secreting airway memory CD4^+^ T-cell responses [33], Th17 responses also contribute to protection against virus infections [21, 34]. These findings suggest that RBD-specific T-cell responses induced by *3Rs-NC+KFDi.n* may enhance protection against these three clusters of hACE2-using sarbecovirus, and potentially offer more conserved cross-reactive protection.

Interestingly, we observed significant differences in infection outcomes and vaccine efficacy between sexes. In the rRsSHC014S challenge model, all five male hACE2 mice in the saline group succumbed between 6 to 8 dpi, whereas mortality in female hACE2 mice was first observed at 8 dpi, with one female mouse surviving at 10 dpi. These observations are consistent with previous studies in hACE2 mice and clinical data that have identified male sex as a risk factor for mortality associated with SARS-CoV-2 infection [22, 35]. Additionally, male hACE2 mice exhibited a significantly lower level of RBD-specific serum IgG responses compared to female hACE2 mice, especially in the *3Rs-NC+KFDi.n* group (**Fig. 5** and supplementary figure 5). We speculate that the lower RBD-specific serum IgG responses and higher risk of mortality in male hACE2 mice may explain the less promising protection against the rRsSHC014S challenge in this group.

In summary, *3R-NC+KFDi.n* induces potent cross-reactive RBD-specific antibodies in both serum and mucosal sites, conferring broad protection against all three sarbecovirus clusters. Furthermore, the vaccine-induced RBD-specific antibodies are long-lasting, providing long-term protection in upper- and lower-respiratory tracts for at least 12 months. Combined with the high safety profile of the mucosal adjuvant KFD, needle-free administration [9], and its ease of manufacturing, *3Rs-NC+KFDi.n* represents a promising mucosal vaccine candidate against all three clusters of hACE2-using sarbecorius. This approach could be extended to develop a comprehensive coronavirus vaccine package targeting non-human ACE2-using sarbecoviruses, human ACE2-using sarbecoviruses, and merbecoviruses.

## MATERIALS AND METHODS

### Mice and ethics

Female 6-8-week-old BALB/c mice were purchased from Beijing Vital River Laboratory Animal Technology Co. Ltd, Beijing, China. The HFH4-hACE2 transgenic mice on C57BL/6 background (kindly provided by Ralph Baric) were propagated and bred at the Laboratory Animal Center. Mice were randomly assigned to groups. All mice were raised in individually ventilated cages (IVCs) under specific pathogen-free (SPF) conditions. The infection experiments on hACE2 transgenic mice were performed in the Animal Biosafety Level 3 (ABSL-3) Laboratory. Animal studies were approved by the Animal Welfare and Ethical Review Committee of WIV, and conducted according to Regulations for the Administration of Affairs Concerning Experimental Animals in China (study number 09202101). The infection experiments on BALB/c mice were performed in the Animal Biosafety Level 3 (ABSL-3) Laboratory, which were approved by the Institutional Animal Care and Use Committee (AUP #WP2022-0044) and the Institutional Biosafety Committee (IBC, Protocol #S0132240E).

### Phylogenetic tree and antigenic distance mapping

All publicly available full genome sarbecoviruses sequences were collected from GenBank or downloaded from GISAID (Supplementary Table 1). Then the MEGA12 program was used with a maximum likelihood algorithm to infer 28 RBD sequences for each sarbecoviruses clade. Principal Coordinate Analysis (PCoA) was then performed on these distance matrices. The first two principal coordinates (PCoA1 and PCoA2), representing the highest variance components, were extracted to generate a 2D scatter plot. Variance explained by each axis was calculated based on corrected relative eigenvalues.

### Vaccine preparation

N-terminal and C-terminal regions of D0-D1 gene of *E. coli* K12 strain MG1655 were linked to construct the KFD gene, cloned into pET-28a plasmid vector (Invitrogen), transformed into *E. coli* BL21 DE3 strain. With 6×His tag in C-terminus, recombinant protein KFD were induced by IPTG plus lactose.

The scaffold NC was designed based on the 3D structure of KFD as previously described [8]. One RBD (aa. 319-527) of SARS-CoV-2 Omicron variant BA.1 (B.1.1.529), N region of scaffold, one RBD (aa. 307-514) of SARS-like virus RsSHC014, C region of scaffold and another RBD (aa. 307-514) of SARS-like virus WIV1 were sequentially linked to construct the 3Rs-NC gene. One RBD (aa. 319-527) of SARS-CoV-2 Omicron variant BA.1 (B.1.1.529), N region of scaffold, one RBD (aa. 319-527) of Delta variant, C region of scaffold and another RBD (aa. 319-527) of Omicron variant were sequentially linked to construct the 3Ro-NC gene [8].

With the presence of signal peptide tPA in the 5’ region and 8×His tag in the 3’ region, RBDs of Omicron variant BA.1 (Ro), RsSHC014 (Rs), WIV1 (Rw), 3Rs-NC, and 3Ro-NC genes were cloned into pcDNA 3.1 plasmid vector, respectively. The recombinant plasmids were transfected into 293F cells in the presence of polyethyleneimine (PEI). Culture supernatants were collected after grown for 72 h at 37℃with 5% CO_2_ and shaking.

These recombinant proteins were purified from by affinity chromatography on a Ni-NTA column (QIAGEN). The residual LPS was removed and determined using the Limulus assay (Associates of Cape Cod) to be less than 0.02 EU/μg protein.

### Vaccination

6-8 weeks old female BALB/c or 10-14 weeks old hACE2 mice were intramuscularly immunized with immunogen plus 200 μg Alum adjuvant (ImjectTM, Thermo Fisher) in 50 ul PBS Imject^TM^ in the lower hind limb or intranasally immunized with 3Rs-NC or 3Rs-NC plus 1 μg KFD in 10 ul PBS three times. Intranasal immunization was performed after anesthesia with pentobarbital sodium (50 mg/kg).

### Samples collection and antibody measurement

Mice sera and mucosal secretions were collected after immunization for analysis of RBD -specific antibody responses as previously described [18, 36]. Briefly, vaginal lavage fluid was collected by aspirating 40µL of PBS into the vaginal tract with 200µL tips. The fluid was blown and aspirated 10 times, washed twice and mixed together. Saliva was procured after carbamylcholine (Sigma) intraperitoneal injection. Serum was obtained by collecting blood samples from the retro-orbital plexus after anesthesia. The mucosal samples were centrifuged at 4000 rpm for 10 minutes before being stored.

All samples were stored at −80°C until they were assayed by ELISA. Alkaline phosphatase-labeled goat anti-mouse IgG, IgA (Southern Biotechnology) polyclonal antibodies and substrate (p-nitrophenyl phosphate; Sigma) were used for detection. ODs were read at 405 nm by an ELISA plate reader (Thermo Labsystems).

### Virus preparation and Neutralization assay

To produce SARS-CoV-2 spike pseudotyped virus, 60 μg of plasmid pNL4-3.luc.RE and 20 μg of different variants of SARS-CoV-2 Spike were co-transfected with PEI into 15cm cell culture dish of HEK293T-cells. The supernatant was harvested 72 h post-transfection, centrifuged at 1000 rpm and stored at –80℃ until use. To assess the neutralizing efficiency of serum, the samples were serially diluted with DMEM supplemented with 10% FBS from 1:10 to 1:2560 in a total volume of 50 μl and then co-incubated for 1h at 37℃with the 20μl of 200 50% tissue culture infectious doses (TCID50) virus of pseudo-typed virus. About 3×10^5^ ACE2-293T-cells in 30 μl complete media were added per well and incubated for 48 h at 37℃ supplied with 5% CO_2_. Luciferase activity was analyzed by the luciferase assay system (Promega).

The SARS-CoV-2 Omicron strain BA.1 (IVCAS6.7600) [8], rRsSHC014S [17], WIV1 [15] were provided by the National Virus Resource Center (Wuhan, China). The virus was propagated and titrated in Cercopithecus aethiops kidney cells (Vero-E6, ATCC CRL-1586). To assess the neutralizing activity of the serum, samples were serially diluted from 1:10 to 1:10240 and incubated with 100 PFU (plaque-forming unit) at 37℃ for 0.5 h. The serum/virus volume ratio was 1:1. The mixtures were then added to Vero-E6 cells and incubated at 37°C for an additional 1 h. The inoculum was removed and the cells were incubated with 0.9% methylcellulose for 5 days at 37℃ supplied with 5% CO_2_.

The 50% neutralization titers (NT_50_) were determined by a four-parameter logistic regression using GraphPad Prism 8.0 (GraphPad Software Inc.) as previously described [16].

### Immunocytes Isolation

For isolation of pulmonary immunocytes in mice, lungs were removed and cut into pieces and digested in 2 mL of HBSS buffer (HyClone) containing 0.5 mg/mL collagenase 1 (GIBCO) and 0.2 mg/mL Dnase I (Sigma-Aldrich) for 45 minutes at 37°C. After digestion, the tissues were minced and pressed through a 70 μm nylon mesh screen, and the resulting single-cell suspensions were collected and washed extensively with cold phosphate buffer saline (PBS). Then the cell pellets were resuspended in 40% Percoll (GE Healthcare) and layered carefully onto 70% Percoll to generate discontinuous Percoll gradients and centrifuged at 2000 rpm for 25 minutes at 22°C. Cells were aspirated in the interface between 40% and 70% Percoll gradients.

For isolation of splenic lymphocytes in mice, spleens were minced and pressed through the 70μm nylon mesh screen. Then lymphocytes were obtained after using Mouse RBC Lysis Buffer (Dakewe).

### ELISPOT

For simultaneous detection IFN-γ, IL-17A, the Mouse IFN-γ/IL-17A FluoroSpot kit (Mabtech) was used. Briefly, 5×10^5^ of lymphocytes from spleen or cervical lymph nodes were seeded into the pre-coated FluoroSpot plate and stimulated with 2 μg/mL SARS-CoV-2 RBD at 37℃, with 5% CO_2_ for 42 hours. The cell culture detection mAbs mix, Fluorophore conjugates mix, and FluoroSpot enhancer were sequentially added. At last, spots were developed by AEC coloring system, and counted by automatic EliSpot System Classic (AID, Germany).

### Mice challenge

For challenge experiments on HFH4-hACE2 mice, the immunized mice were intranasally inoculated with 6× 10^4^ TCID_50_ of SARS-CoV-2 Omicron strain BA.1 (IVCAS6.7600) or SARS like strain WIV1 in 30 μL under avertin (250 mg/kg) anesthesia. At 3 days post-infection, after euthanized, the lung and turbinate tissues of mice were harvested.

For rRsSHC014S challenge experiments, the immunized HFH4-hACE2 mice were intranasally inoculated with 1× 10^5^ TCID_50_ of rRsSHC014S in 40 μL under avertin (250 mg/kg) anesthesia. Weight and death of mice were recorded every day until at 10 days post-infection.

For challenge experiments on BALB/c mice, the immunized mice were intranasally inoculated with 4.35 × 10^4^ TCID50 of Omicron strain BA.1 (BA1-HB0000428) in 50 μl under avertin (250 mg/kg) anesthesia. At 4 dpi, after euthanized, the lung and turbinate tissue.

### RNA extraction and RT-qPCR

For testing sarbecovirus in the tissues of infected HFH4-hACE2 mice, tissues were homogenized in DMEM and viral RNA was extracted using the FineQuick Viral DNA/RNA Kit-automated version coupled with an automated Purifier Modesty instrument (Genfine Biotech, Beijing, China). Viral Np gene was detected using the HiScript II One Step qRT-PCR SYBR Green Kit (Applied Biosystems, USA). Viral sequence copies were determined by comparing the Ct values to a standard curve derived from a cDNA clone.

For testing virus in the tissues of SARS-CoV-2 Omicron strain BA.1 (BA1-HB0000428) challenged BALB/c mice, tissues were homogenized in DMEM and 0.1 mL of clarified homogenate supernatant fluid was mixed with 0.4 mL of TRIzol LS Reagent (Invitrogen), and the total RNA was extracted according to the manufacturer’s instructions. Next, first-strand cDNA was synthesized using a PrimeScript RT kit (Takara), and real-time quantitative PCR (RT-qPCR) was used to detect the presence of SARS-CoV-2 distribution. Copies of viral E gene and the internal reference GAPDH gene were determined by comparing the Ct values to a standard curve derived from a cDNA clone.

### Statistical analysis

Means, SD, neutralizing titers and correlations were calculated using GraphPad Prism 8.0 software. Test for significance was applied as indicated in the legend of each figure. Except otherwise indicated, statistical analysis was performed using one-way ANOVA followed by Dunnett’s multiple comparison test. To analyze the differences between two groups, unpaired two-tailed Student’s *t*-test was used for normally distributed data with homogeneous variance, and the Mann-Whitney U test was used for nonnormally distributed data. Simple linear regression was used for correlation analysis. Analyses were performed using GraphPad Prism 8.0 software. Significance values are indicated as *p < 0.05, **p < 0.01, and ***p < 0.001; ns, nonsignificant. p < 0.05 was considered significant.

## List of Supplementary Materials

supplementary table 1

supplementary figure 1 to 7

## Acknowledgments

We especially thank Tao Du and Jin Xiong for their technical assistance and equipment support in the biosafety level 3 (BSL-3) facility. We also thank Xuefang An and Fan Zhang (core facility, Wuhan Institute of Virology, CAS) for their technical support and kind help in animal experiments.

## Funding

This work was supported in whole or in part by the Shanghai Science and Technology Innovation Action Plan (Grant number: 22Y11901000 to Q.W.), National Key R&D program of China (Grant number: 2021YFC2302602 to J.Y.), National Natural Science Foundation of China (Grant number: 82341041 to H.Y. and 32100749 to M.L.), Shenzhen Medical Research Fund (Grant number: B2302044 to Y.-Q.C.), Shenzhen Science and Technology Program (Grant number: RCJC20210706092009004, KOTD20200820145822023, ZDSYS20230626091203007 to Y.-Q.C.).

## Author contributions

H.Y., K.L, Y.C, L.Z J.Y. Y.-Q.C. and Q.W conceptualized and supervised the study. J.Y., H.L., L.L., Y.-Q.C. and L.Z. developed Methodology. L.L., H.L., M.L., X.L., M.X., Y.H., M-Q.L., Z.H. and Z.Z. performed the investigation. J.Y., W.S. and S.L. visualized the data. J.Y., M.L., H.L., L.L., L.Z., Y.-Q.C. and Q.W. wrote the original draft. All authors reviewed and edited the manuscript.

## Competing interests

The authors declare that they have no competing interests.

## Data and materials availability

All data associated with this study are present in the paper or the Supplementary Materials.

## Competing interests

The authors declare that they have no competing interests.

## Notes

### Competing Interest Statement

The authors have declared no competing interest.

