## Supplemental Table 1 and Supplemental Figures 1-7 for "A Novel ‘Three-in-one’ Mucosal Vaccine Elicits Broad Protection against Three Distinct Clusters of ACE2-using Sarbecoviruseses"

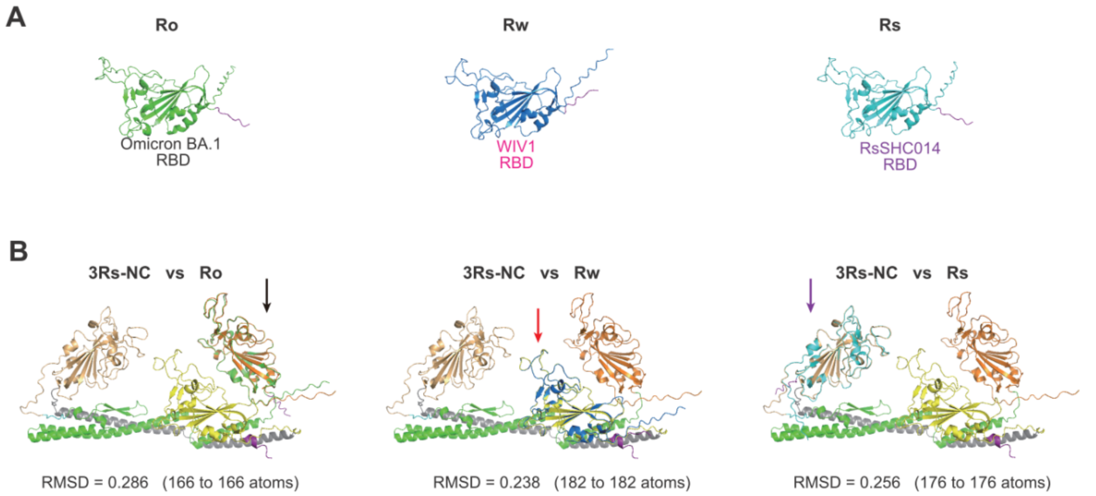


**Supplementary Fig. 1. RBD domains within 3Rs-NC have similar 3D structures with monomeric RBDs (**Related to Fig.1**).** (**A**) 3D structure of the RBD monomer derived from SARS-CoV-2 Omicron BA.1 strain RBD (Ro), WIV1 RBD (Rw), and RsRSHC014 RBD (Rs) predicted by AlphaFold 3. (**B**) Pairwise alignments of predicted 3D structures of 3Rs-NC with Ro, Rw, Rs via Pymol, respectively. Arrows indicated the aligned sites.

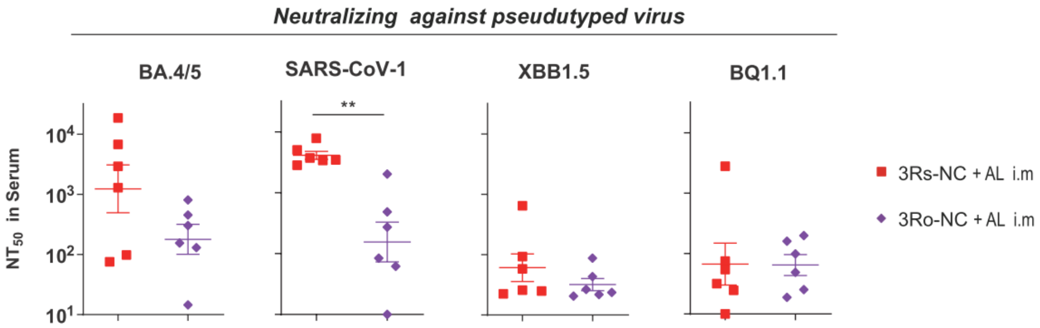


**Supplementary Fig. 2. Neutralizing antibody responses in serum after the 3rd immunization against pseudotyped SARS-CoV-1 and SARS-CoV-2 variants** (Related to Fig.2)**.** Data are represented as mean ± SEM. Groups were compared using an unpaired *t* test. **p < 0.01.

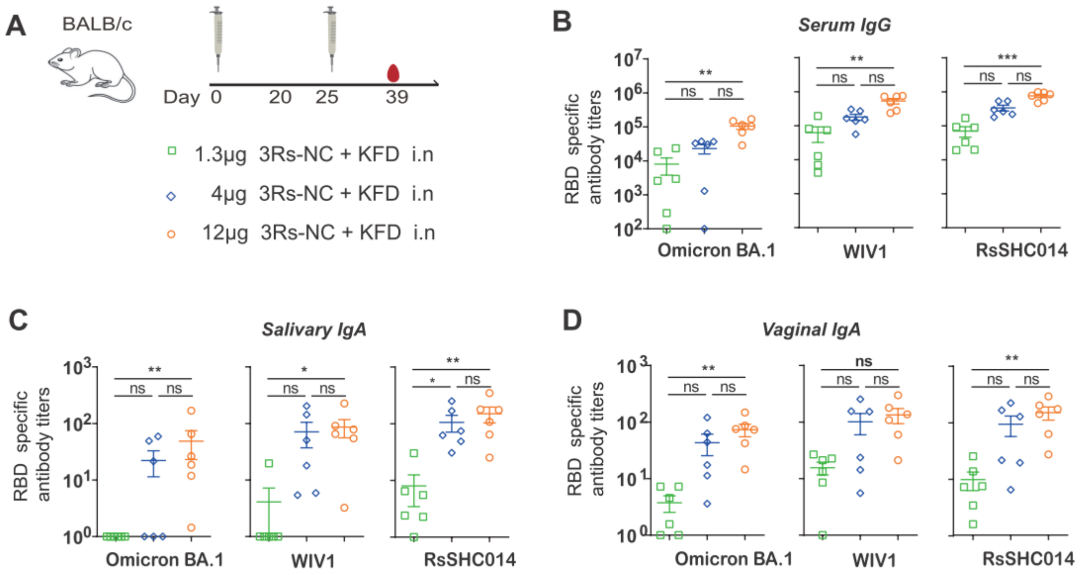


**Supplementary Fig. 3. Antibody responses by the intranasal immunization of different dosages of 3Rs-NC adjuvanted with KFD in BALB/c mice (**Related to Fig. 3**).** (**A**) Diagram scheme of immunization and sampling (n=6 female mice per group). (**B**) RBD-specific IgG responses in serum after the 3$^{rd}$ immunization. (**C** and **D**) RBD-specific mucosal IgA responses after the 3$^{rd}$ immunization in saliva (**C**) and vaginal lavage fluid (**D**). Data are represented as mean ± SEM. Groups were compared using one-way ANOVA. *p < 0.05; **p < 0.01; ***p < 0.001; ns, nonsignificant.

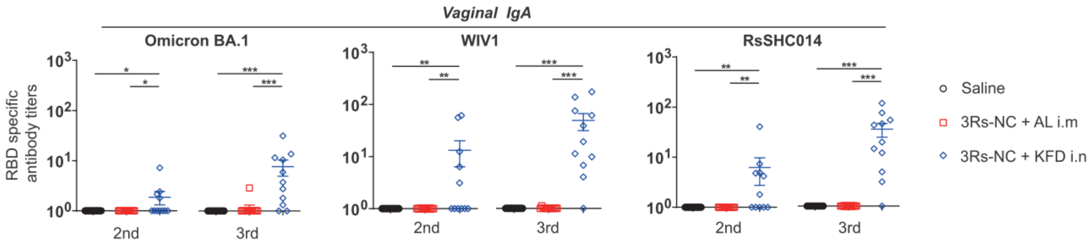


**Supplementary Fig. 4. RBD-specific vaginal IgA after the 2nd and 3rd immunization of 3Rs-NC with adjuvants in HFH4-hACE2 mice** (Related to Fig.4)**.** RBD-specific vaginal IgA after the 2nd and 3rd immunizations. Groups were compared using one-way ANOVA. *p < 0.05; **p < 0.01; ***p < 0.001.

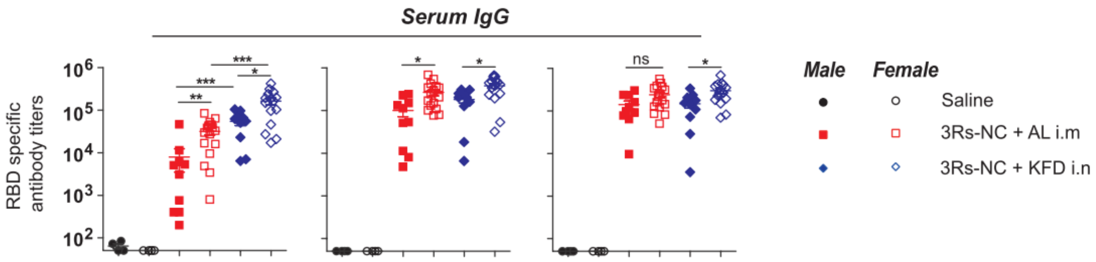


**Supplementary Fig. 5. RBD-specific serum IgG after the 3rd immunization of 3Rs-NC with adjuvants in HFH4-hACE2 mice (**Related to Fig.4 and Fig.5**).** Groups were compared using one-way ANOVA. *$p < 0.05$; **$p < 0.01$; ***$p < 0.001$; ns, nonsignificant.

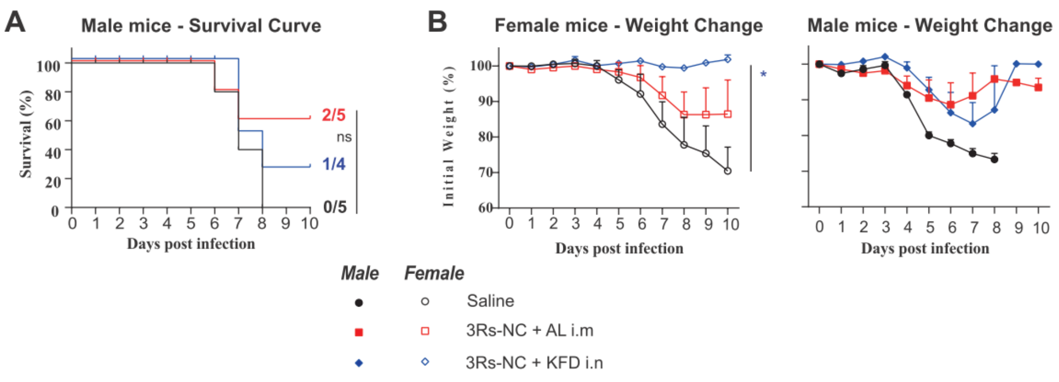


**Supplementary Fig. 6. Protection of 3Rs-NC immunization against virus rRsSHC014S in male and female HFH4-hACE2 mice (**Related to Fig.5**). (A**) Survival curves of male mice within 10 days post-infection. (**B**) Weight changes of female and male mice within 10 days post infection. Groups were compared using one-way ANOVA. *$p < 0.05$.

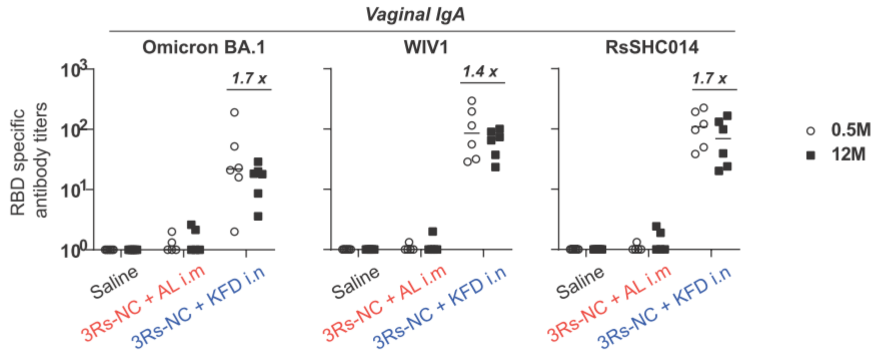


**Supplementary Fig. 7. Long-lasting RBD-specific antibody in vaginal lavage fluid of 3Rs-NC immunized mice (**Related to Fig.6**).** RBD-specific vaginal IgA at 0.5 months and 12 months after the 3rd immunization.

**Supplementary Table 1** List of sequences and accession numbers used in this study.

| Name | Accession number |
|---|---|
| WIV16 | KT444582 |
| CUHK-W1 | AY278554 |
| SZ/6103 | AY515512 |
| GD03T0013 | AY525636 |
| RsSHC014 | KC881005 |
| Rs3367 | KC881006 |
| WIV1 | KF367457 |
| BtRs-BetaCoV/YN2018B | MK211376 |
| SARS-CoV-2 (Prototype) | MN908947 |
| RaTG13 | MN996532 |
| bat-SL-CoVZC45 | MG772933 |
| bat-SL-CoVZXC21 | MG772934 |
| BtRl-BetaCoV/SC2018 | MK211374 |
| Rf1/2004 | DQ412042 |
| Bat CoV 273/2005 | DQ648856 |
| BtRs-BetaCoV/YN2018D | MK211378 |
| HKU3-1 | DQ022305 |
| Rm1/2004 | DQ412043 |
| BtCoV/279/2005 | DQ648857 |
| Rp3/2004 | DQ071615 |
| BtRs-BetaCoV/YN2018A | MK211375 |
| BtRs-BetaCoV/YN2018C | MK211377 |
| SARS-CoV-2 (Beta) | EPI_ISL_736940 |
| SARS-CoV-2 (Delta) | OK091006.1 |
| SARS-CoV-2 (BA.4/5) | EPI_ISL_12029894 |
| Pangolin/GD/1 | EPI_ISL_410721 |
| BtKY72 | APO40579.1 |
| BM48–31/BGR/2008 | GU190215 |
